# Loss-of-function mutations in novel triacylglycerol lipase genes are associated with low rancidity in pearl millet flour

**DOI:** 10.1101/2022.04.02.486827

**Authors:** Rasika Rajendra Aher, Palakolanu Sudhakar Reddy, Rupam Kumar Bhunia, Kayla S. Flyckt, Aishwarya R Shankhapal, Rabishankar Ojha, John D. Everard, Laura L. Wayne, Brian M. Ruddy, Benjamin Deonovic, Shashi K. Gupta, Kiran K. Sharma, Pooja Bhatnagar-Mathur

## Abstract

Commercialization and utilization of pearl millet (*Pennisetum glaucum* L.) by consumers and processing industry is constrained due to rapid onset of rancidity in its milled flour. We studied the underlying biochemical and molecular mechanisms to flour rancidity in contrasting inbreds under 21-day accelerated storage. Rapid TAG decrease was accompanied by FFA increase in high rancidity genotype compared to the low rancidity line, that maintained lower FFA and high TAG levels, besides lower headspace aldehydes. DNA sequence polymorphisms observed in two lipase genes revealed loss-of-function mutations that were functionally confirmed in yeast system. We outline a direct mechanism for mutations in these key TAG lipases in pearl millet and the protection of TAG and fatty acids from hydrolytic and oxidative rancidity respectively,. Natural variation in the *PgTAGLip1* and *PgTAGLip2* genes may be selected through marker assisted breeding or by precision genetics methods to develop hybrids with improved flour shelf life.

## 1. Introduction

Pearl millet (*Pennisetum glaucum* L.) is the principal staple for millions of people in arid and semi-arid regions of Asia and Africa, such as India and Nigeria. It is primarily grown in the driest regions due to its ability to tolerate and thrive under both continuous and erratic drought. This C4 plant requires low inputs when compared with other staples such as rice and wheat, effectively reduces atmospheric CO_2_, has high water use efficiency, and keeps drylands productive, ensuring food and nutritional security of smallholder farming communities. A powerhouse of nutrients like protein, minerals, vitamins, phytochemicals and antioxidants, pearl millet is gluten-free and contains high levels of polyphenols and other biologically advantageous compounds, accordingly has been categorized as a “Smart food” (Tako et al., 2015). Its beneficial health properties such as lowering of fat absorption in the intestines and spike-free sugar release (low glycaemic index) helps reducing high blood pressure, heart disease, and diabetes (Varsha and Narayana, 2017).

Despite its nutritional profile and advantage over other cereals, pearl millet has remained unpopular due to short storage life of the milled flour (Goswami et al., 2020). The rapid development of off flavours and taste within 5-7 days of milling poses a major hindrance for wider consumer acceptance, creates post-harvest waste, and causes drudgery to consumers, particularly women who are traditionally involved in food preparation. Over the last decade, pearl millet consumption has declined in urban areas and in commercial outlets, leading to lower incentives for its cultivation by smallholder farmers. To address rancidity, efforts have been made to develop post-harvest processing and pre-milling techniques to inactivate the biological components that lead to accelerated rancidity. However, these mechanical and physiochemical techniques have not been very effective, and in fact have been shown to negatively influence the nutritional quality of both the micro- and macro-nutrients (Goyal et al., 2015). Therefore, to revive the importance of this nutri-cereal in dryland agriculture, further understanding of the biological processes that lead to the development of rancidity in pearl millet is urgently needed.

Pearl millet grain has a larger germ layer than other cereals, apart from maize (Taylor, 2004), and has a higher lipid content (5-7%), characterised by high levels of unsaturated fatty acids (Sharma et al., 2015). During milling, the bran and germ layer rupture and releases endogenous lipases that commence the hydrolysis of stored lipids and release of free fatty acids (FFAs). Lipases are more thermostable than lipoxygenases and in low moisture conditions, show higher enzyme activity. Hence, inhibiting the initial formation of FFAs is critical for controlling seed lipid oxidation. While lipases are reported to play a primary role in the susceptibility of pearl millet to post-milling rancidity (Kumar et al., 2021), the underlying biochemical and molecular mechanisms have not yet been studied in detail, and to our knowledge the specific lipase(s) responsible have not been confirmed.

Here, we demonstrate that flour from a pearl millet flour characterized as having low rancidity is more resistant to TAG degradation and lipid oxidation than that milled from high rancidity germplasm. Two TAG lipase genes (*PgTAGLip1 and PgTAGLip2*) in the low rancid-pearl millet line contained polymorphisms that rendered the enzymes non-functional. These key TAG lipases may be used to develop seeds with greater post-milling shelf life, without affecting the beneficial nutritional parameters.

## 2. Materials and Methods

### 2.1. Plant material and growth

Initial screening experiments were performed on a representative panel of 12 inbred and inbred germplasm association lines (PMiGAP) of pearl millet (*Pennisetum glaucum* L.) developed by the millet breeding program at ICRISAT (Table S1). These lines were grown at the Patancheru campus in India and harvested during the rainy season of 2018 and 2019, and analysed in 2019 and 2021, respectively.

### 2.2. Flour milling and storage conditions

Seed was stored in vacuum packed pouches in a cold room prior to analysis. Thirty gram of seed samples of each line were ground to fine powders in a custom-built burr-mill grinder or in a Cyclotec (Foss) grinding mill. The ground flours were spread into evenly distributed layers in lidless food grade trays (4 oz/118 mL) under accelerated storage conditions in incubator chambers at a temperature of 35 +/− 0.1 °C and 75 +/− 3 % relative humidity. Sub-samples of the flour were collected at time points for the biochemical determination acid value (AV), triacylglycerol (TAG) break down, FFA generation, and volatiles.

### 2.3. Acid Value

Total crude fat was extracted in a Soxhlet apparatus from 5 g flour using the standard method (AOAC, 1990) with few modifications. Briefly, 5 g of flour was taken in a cellulose thimble which was suspended in pre-weighed extraction beaker containing 100 mL petroleum ether (boiling point 60-80 °C) and kept on a hot plate at 180 °C until the sample started to boil. For the complete extraction of fat, the process was carried out for at least 90 min. Following removal of the solvent, the extracted oil was titrated against 0.1 N KOH using phenolphthalein as indicator and the end point recorded. The acid value was calculated using the formula:

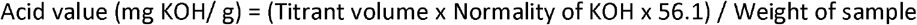

### 2.4. Quantification of Lipids by GC-FID and HPLC-ELSD

The lipid extractions were based on the method of Bligh and Dyer (1959). Briefly, lipids were extracted using 3 mL of 1:2 chloroform: methanol by shaking at room temperature for 15 min. Phase separation was induced by the addition of 1 mL chloroform and 1.8 mL water. The lipid extracts were subsequently washed with 0.5 mL of 1 M KCl. The lower lipid layer was collected, and samples re-extracted twice with 2 mL chloroform. The lipid extracts were subsequently washed with 0.5 mL of 1 M KCl, filtered, and dried under a stream of nitrogen gas. In the 2021 experiment, the extraction method was modified to include an initial step to quench lipase activity by treatment with isopropanol heated to 75° C for 15 min and 0.01% BHT was added to all reagents.

For lipid quantitation by GC-FID, 0.1 mg of 17:0 TAG was added to each sample as an internal standard. For quantitation of total fatty acids, acid catalysis was performed by adding 1 mL of 5% sulfuric acid in methanol followed by heating at 90°C for 1 h. Phase separation was induced by the addition of 1 mL of 1M NaCl and 1 mL heptane. The upper heptane layer was transferred to a sample vial for analysis. FAME separation was performed on a GC system (Agilent 7890A) with an OmegaWax 320 column (Supelco) followed by FID analysis. The GC oven temperature was set at a starting temperature of 190 °C, then increased to 240 °C @ 5 °C/min, with a total run time of 10 min. Neutral lipids were quantified by HPLC equipped with an ELSD (Evaporative Light Scattering Detector) using a cyanopropyl column (Luna 5 μm CN 100Å 250 x 4.6 mm; Phenomenex) with hexane as mobile phase A and methyl tertiary-butyl ether (MTBE):isopropanol (95:5 v/v) plus 0.2% acetic acid as mobile phase B, with a gradient of 0% to 100% B, and re-equilibration of the column to 0% B. Standard curves of tri-C18:2 TAG and 18:2 FFA were run with each sample set to quantify total TAG and FFA as a percent of dry weight (% DW).

### 2.5. Analysis of volatiles by Solid Phase Micro-Extraction-GC-MS

Samples for each pearl millet line were subject to Solid Phase Micro-Extraction Gas Chromatography with Electron Impact Ionization Mass Spectrometry (SPME-GC-EI-MS) after 21 d of accelerated storage from the 2019 experiment. Triplicate flour samples (0.5000 +/− 0.0001 g) of each line were weighed into 20 mL amber headspace vials that were brought to room temperature before incubation at 35 °C for 1 h. Analysis was performed on an Agilent 7890 GC system equipped with an Agilent 5977B detector and a Gerstel Multi-Purpose Sampler (MPS Robotic; Gerstel) with automated SPME sampling capability. MS data files were pre-processed and analysed with Genedata Expressionist software tools. Details on chromatography hardware parameters, software for processing and statistics is provided in Supplementary Material & Methods.

### 2.6. RNA isolation, cDNA synthesis and quantitative real time-PCR (qRT-PCR) analysis

Pearl millet tissue samples were collected at 0 h, 6 h, 12 h, and 24 h and used for total RNA extraction using RNeasy Plant Mini kit (Qiagen, Germany), and analyzed quantitatively and qualitatively using NanoVue plus spectrophotometer (GE health care, USA). One and a half μg of RNA was used for cDNA synthesis using Superscript III (Invitrogen) and samples diluted 1:10 times were used as template. Quantitative RT-PCR was carried out in 96-well optical reaction plates with a total reaction volume of 10 μL containing 0.5 μM of each primer (0.8 μL), cDNA (1.0 μL), Sensi Master Mix (2X) and dH_2_O added up to 3.2 μL. The qRT-PCR primers were designed using Primer3 software (v.0.4.0) with GC content of 40–60%, a T_m_ of 60–62 °C, and primer length 20-25 nucleotides (Table S3), for an expected product size of 90–180 bp. The qRT-PCR reactions were carried out using a standard thermal profile: 95 °C for 10s, 40 cycles of 15s at 95 °C, 15s at 61 °C with fluorescent signal recording and 15 seconds at 72 °C. After the 40^th^ cycle, amplicon dissociation curves were measured by heating at 58 to 95 °C with fluorescence measured within 20 min. All qRT-PCR data were obtained from three biological replicates with three technical replicates. Normalized expression was calculated with qBase+ software (Schmidt and Delaney, 2010) with reference genes *eukaryotic initiation factor4α* (*PgEIF4α*) and *Malate dehydrogenase (PgMDH*) (Reddy *et al*., 2015). List of all primers used for all molecular studies is given in Table S2.

Gene expression analysis was carried out using vegetative tissues (leaf, shoot and roots), seed developmental stages (embryo, milky seed, immature seed), and harvested grain flour under accelerated storage for 24 h. Lipid reserve mobilization studies were conducted on filter paper germinated seedlings sampled at regular intervals during 0-7 days after imbibition (DAI).

### 2.7. Sequence analysis of the pearl millet TAG Lipase genes

Rice lipase protein sequences were used as the query and searched in the latest version of the pearl millet genome (Varshney et al., 2017). Following removal of all duplicate and redundant sequences, the remaining samples were analysed for prediction of domains and motifs using the ExPASy PROSITE tool. Multiple sequence alignment was carried out with full-length protein sequences of PgTAGLips with the ClustalW programme of MacVector assembly programme (V17.1). Evolutionary relationship within lipases of pearl millet were investigated using the neighbour-joining algorithm of MacVector Software and the subcellular localization of proteins was predicted using the WoLF PSORT tool.

### 2.8. Molecular modelling and docking

Molecular models of PgTAGLips were generated using homology modelling server SWISS MODEL (Waterhouse et al., 2018) by utilizing the known structure of the template protein. Ramachandran plots were generated using PROCHECK (Laskowski et al., 1993) for validation of structures. Three dimensional structures of triglycerides and ester ligands substrate were retrieved using the PubChem database. Autodock 4.2 (Morris et al., 2009) was used for docking the triglycerides substrates to the lipase gene structures from the simulation. A grid box was prepared with dimensions 30X30X30 and centred on the ATP binding site with grid spacing of 0.478 Å. Ten docking runs were performed for each substrate with a population size of 50 and 15,00,000 energy evaluation. All other algorithm parameters used the default setting. The docked poses for each ligand were clustered with an RMSD tolerance of 1.5 Å. Predicting the possible protein-ligand interactions and the final pose of the substrate was selected based on its docking score.

### 2.9. Gene cloning

Selected PgTAGLips were PCR amplified using cDNA as a template and cloned in pCR8/GW/TOPO TA Cloning vector (Invitrogen). All sequence confirmations were carried out with at least four to five colonies from each genotype. The vector backbone was trimmed and sequences analysed and aligned across genotypes using ClustalW software to identify sequence variabilities.

### 2.10. Yeast transformation

The cDNAs of PgTAGLip genes from selected genotypes were cloned in pYES2 vector under the control of galactose-inducible promoter and URA3 as a yeast selection marker. The yeast triple lipase mutant *Δtgl3Δtgl4Δtgl5* (ΔTGL) (Klein *et al*., 2016) was used for TAG lipase heterologous characterization. All pYES2-PgTAGLip variants were transformed into ΔTGL according to the Ojha *et al*. (2021). Positive yeast transformants were selected on minimal synthetic defined (SD) base (Takara) media supplemented with complete supplement mixture without uracil (−Ura) (Himedia, India) and further identified by colony PCR using conditions described above. Selected single yeast colonies were transferred into dropout base growth medium (Himedia, India) with 2% galactose and 1% raffinose, and incubated at 30 °C for 24 h for lipid and FACS analysis.

### 2.11. BODIPY^493/503^ staining and yeast lipid degradation analysis

Yeast ΔTGL cells transformed with different PgTAGLip variants were stained with BODIPY^493/503^ dye (4,4-Difluoro-1,3,5,7,8-Pentamethyl-4-Bora-3a,4a-Diaza-s-Indacene, Invitrogen) and used to quantify lipid degradation as an indication of lipase gene function. BODIPY dye was used to label the yeast TAGs according to Bhunia et al. (2021). In brief, 2 μl of 1 mg/mL BODIPY stock solution (prepared in DMSO) was added to 1 mL yeast culture carrying different lipase variants and the samples were kept at 28 °C for 20 min. Following incubation period, yeast cells were centrifuged at 10,000 rpm for 10 min and the cell pellets washed thrice with PBS buffer and suspended in 1 mL of PBS buffer. Yeast cells were further diluted 1:10 in PBS buffer. Overnight cultures of yeast cells expressing TAG lipase were diluted into fresh medium containing cerulenin (10 μg/mL) and terbinafane (30 μg/mL) to block the *de novo* fatty acid biosynthesis and collected at 2 h, 4 h and 6 h intervals. Finally, BODIPY dye was used to label the yeast cells for flow cytometry analysis (Bhunia et al., 2021). Labelled yeast cells collected at different intervals were diluted (1:10) in PBS buffer and used to measure the fluorescence intensity in a flow cytometer. The excitation and emission wavelengths were set at 493 nm and 503 nm, respectively.

### 2.12. Flow cytometry analysis

Flow cytometry analysis was performed using high-speed flow cytometer, BD FACSAria Fusion (Becton Dickinson) to measure the uptake of yeast lipids according to Bhunia et al. (2021). Every flow cytometry event was measured using side scatter (SSC) and LB fluorescence using FITC (530⍰±I5 nm excited at 488 nm) filters. The mean fluorescence intensity values were analysed using BD FACSDiva 8.0.1. The flow cytometer settings of all channels remained the same for all yeast cell sorting procedures.

### 2.13. Statistical analysis

Estimation of all chemical/biochemical parameters were carried in triplicates and analysed using Genstat. Several Log-linear hierarchical mixed models for TAG and FFA were fit using the brms software package (Bürkner, 2017) in the R programming language. The model with the lowest leave-one-out information criteria (Vehtari et al., 2017) was chosen as the final model for inference. For more information see supplementary materials.

## 3. Results

### 3.1 Lipid degradation and off-flavour volatiles are increased in high rancidity pearl millet lines

To examine the rancidification of pearl millet flour, acid values (AV) were determined in the oil extracted from flours of selected pearl millet lines subjected to accelerated storage conditions. Crude fat contents of the flours ranged between 4.2 and 7.2 wt% at the start of the experiment (day 0). The AV of freshly ground flours at day 0 varied between 5.15 to 13.6 mg KOH/g among the genotypes. Under accelerated storage conditions, AV increased significantly in all lines (Figure 1a), although the rates varied substantially between genotypes. For instance, while several lines (e.g., I6 and P19) had AV approaching 100 mg KOH/g after 14 d of accelerated aging, others (e.g., I2, I3, and I8) showed significantly lower AV (< 67.33 mg KOH/g). Based on these results, a subset of inbred lines with early onset of rancidity indicators were selected and re-evaluated in a 21 d experiment (Figure 1b). The I3 inbred line had lower AV, while the I5 and I7 lines had higher AV indicative of greater FFA hydrolysis. A sensory panel also characterised the I3 line as less rancid (e.g., no odour, dry flour) and the I5 and I7 lines as more rancid (e.g., pungent, bad smell) after 21 d of accelerated aging (Table S3).

**Figure 1.**
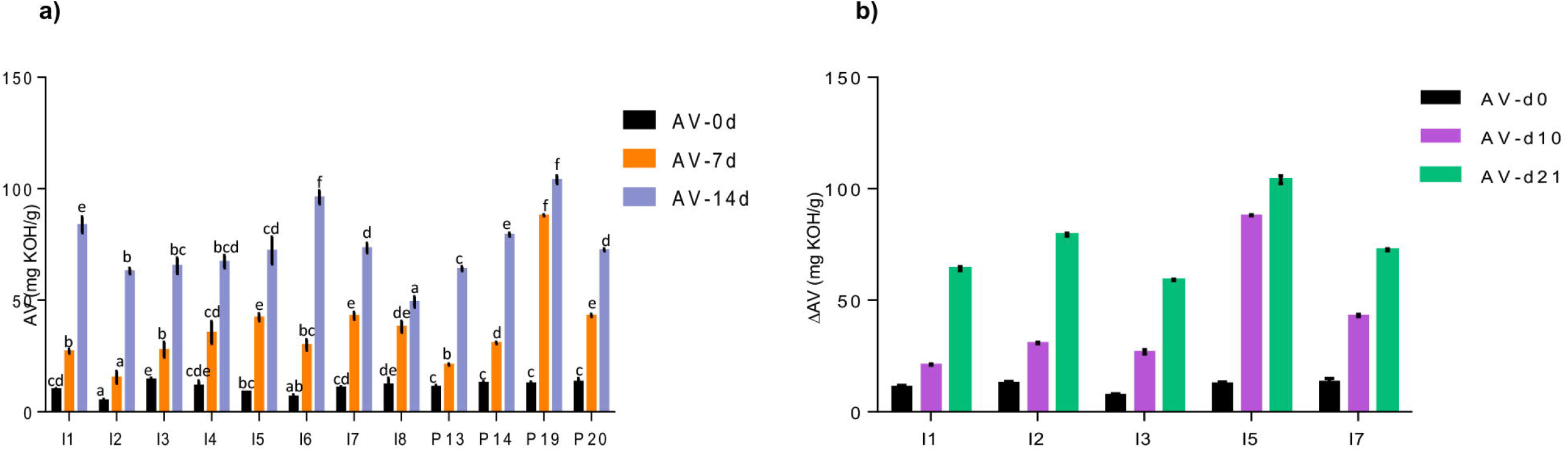
(a) Comparison of Acid value (AV) of flour of 12 pearl millet inbred lines during 14-day accelerated storage conditions. (b) Changes in the AV in the flour of selected contrasting lines on day 10 and 21. The data is represented as mean ± standard error of three biological replicates. Variables showing F-value less than 0.05 were significant. Also, the means displaying nonmatching lowercase letters were significantly different at p ≤0.05 by Duncan test.

The lipids from I3, I5, and I7 flours were analysed to further characterise lipid hydrolysis. All three inbred lines had similar fatty acid profiles dominated by unsaturated linoleic and oleic acids (>75%), which are prone (particularly linoleic acid) to primary and secondary oxidation (Figure 2a). Additionally, pairwise comparisons using two sample Mann-Whitney tests across all fatty acids profiled showed no significant differences, suggesting that fatty acid composition is likely not the cause of differences in the propensity of the lines to develop rancidity. Similarly, total fatty acid measurements showed no significant differences between the lines (Kruskal-Wallis test and pairwise Mann-Whitney tests) in both experiment sets (2019 and 2021), and no significant differences in total fatty acid measurements were observed between the two experiments. Overall, across both experiments, the I7 line had the lowest average total fatty acid content at 3.88%, whereas I3 and I5 on an average comprised of 5.14% and 6.42% total fatty acids, respectively (Figure 2b).

**Figure 2:**
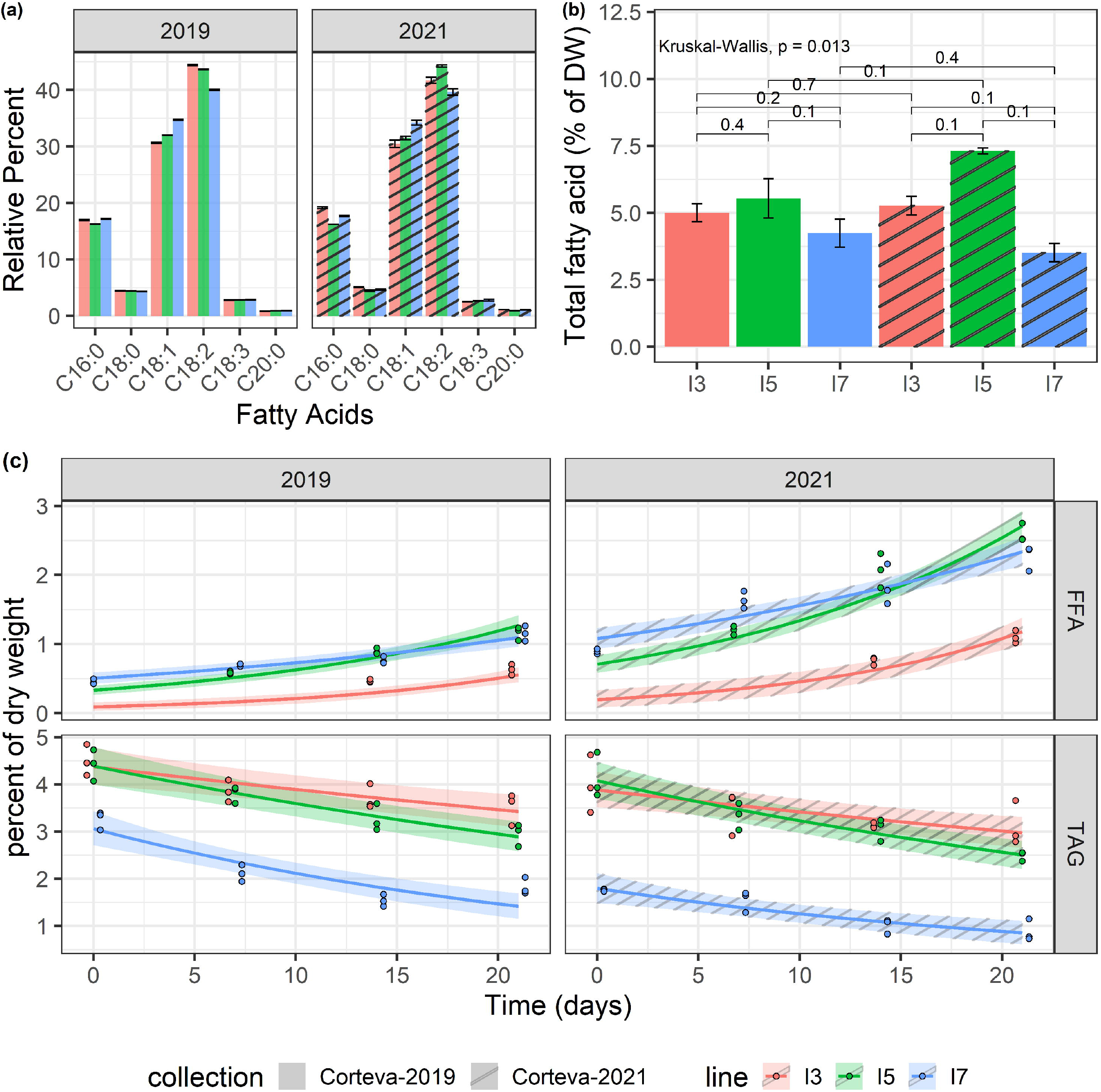
Lipid catabolism in milled flour under accelerated-ageing conditions. (a) Fatty acid profiles obtained by GC-FID. (b) Total fatty acid quantitation by GC-FID, average over three replicates. Each brace corresponds to a Mann-Whitney test with the resulting p-value indicated above the brace. (c) TAG and FFA levels measured by HPLC-ELSD in two datasets (2019 and 2021). Log-linear hierarchical mixed models were utilized to assess the differences in TAG and FFA levels of these three inbred lines. Points represent observed data, lines represent posterior mean of the predictive distribution, and the ribbon corresponds to the 95% credible interval of the posterior mean predictive distribution.

Two independent 21-day accelerated-aging (under high temperature and high humidity) experiments were performed to determine differences in lipid catabolism for the I3, I5 and I7 lines in 2019 (material grown in 2018) and 2021 (material grown in 2019). The TAG and FFA levels were measured via HPLC from subsamples taken at each time point. For all lines, TAG levels showed a decreasing trend overtime and a corresponding increasing trend in FFAs (Figure 2C). Pairwise posterior median differences on the log scale (averaged across the datasets) were computed to assess the significance of the differences between these three lines (Tables S4). The TAG values of the I7 line were significantly lower than the other two lines at all time points. The TAG values of the I3 and I5 line were not significantly different until day 14. All pairwise FFA levels were found to be significantly different except from the I5 and I7 lines at day 14. The I3 line maintained the lowest levels of FFA throughout the experiments. Interestingly, the I7 line exhibited elevated FFA amounts at the onset of the time course across both datasets; it is unknown if I7 seed has elevated FFA pre-milling. Another 2021 dataset, using TLC/ GC-MS, showed similar trends with increasing FFA especially in I7 and decreasing TAG levels over the time course (Figure S1); however, this experiment was not quantitative and there was most likely sample loss. These results suggest that flour from I7 line is more prone to TAG degradation during storage at high temperature and high humidity.

To confirm that the hydrolysis of TAG and the release of FFAs in flours is directly associated with the development of rancidity, the volatile chemicals released from the flours after accelerated storage were analysed using SPME-GC-EI-MS. Principal component analysis of all auto-scaled components indicated that each line was distinct (Figure 3a), however, line I3 appeared to be markedly different from the lines I5 and I7 due to the presence of aliphatic aldehydes and other known markers of lipid oxidation (Figure 3; Table S5). These data clearly indicate differential oxidation of fatty acids in the tested lines. From over 3,000 detected features in this analysis, four of the top 15 most statistically significant metabolites were aldehydes (Figure 3) which are known secondary oxidative products of unsaturated lipids. The I5 and I7 lines had significantly higher levels of hexanal, octanal, nonanal, and benzenacetaldehyde compared to the line I3 in the headspace above flours exposed to 21 days of accelerated aging (Figure 3b, c). Since the fatty acid profiles for the three lines were similar (Figure 2A), the lines I5 and I7 seem to have greater susceptibility to rancidity than the I3 line due to higher amounts of FFAs and the oxidation of these FFAs.

**Figure 3.**
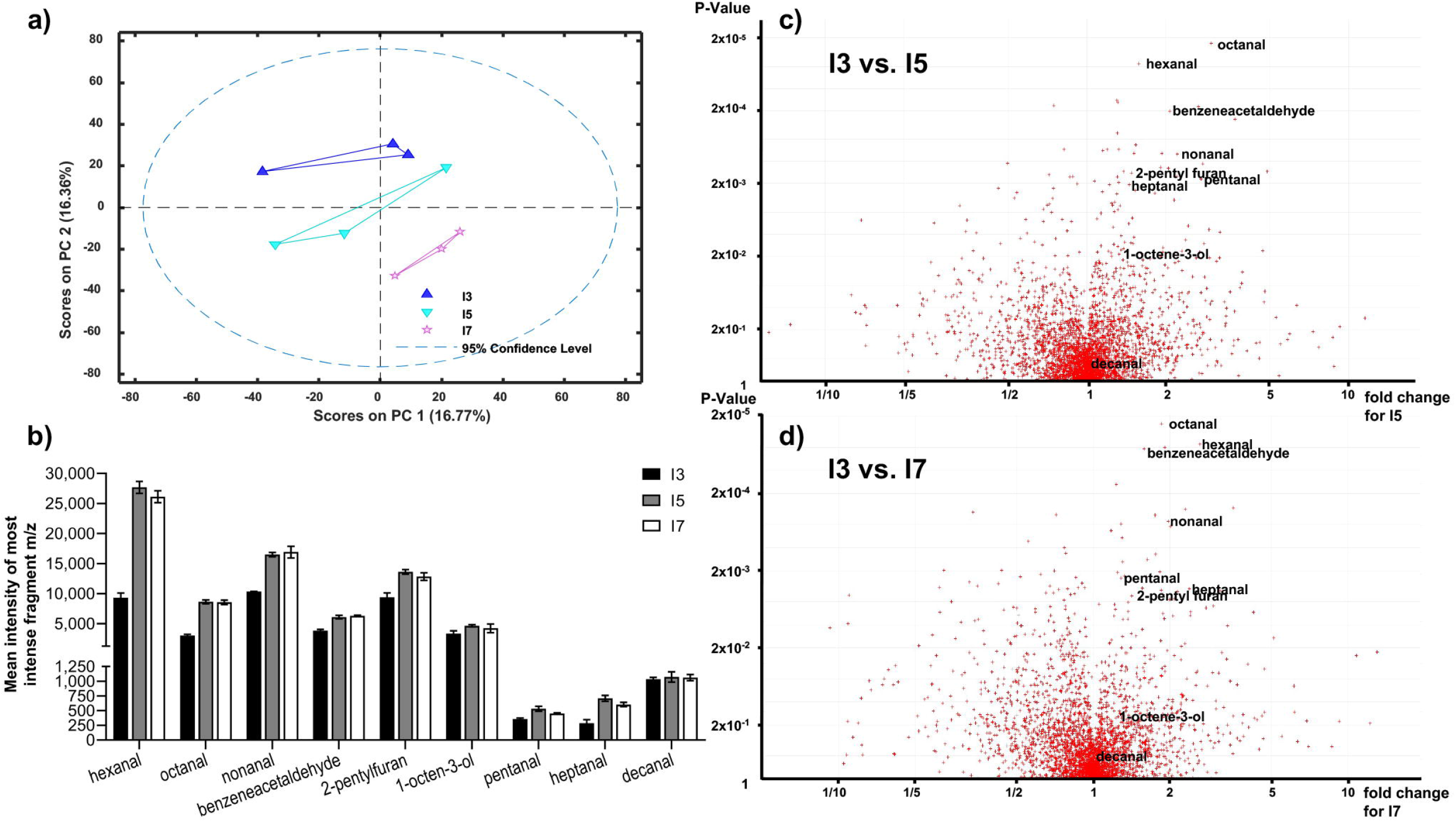
Volatile analysis of pearl millet I3, I5 and I7 lines using Solid Phase Micro-Extraction Electron Impact Ionization Gas Chromatography Mass Spectrometry (SPME-EI-GC-MS). (a) First and second principal components for processed auto-scaled datasets for the I3, I5 and I7 lines (blue triangles, aquamarine inverted triangles and pink stars, respectively). A 95% confidence range is indicated by dashed line. (b) Most abundant fragment m/z total peak intensity bar charts for lowest p-value aldehydes (C_5_-C_10_) for the I3, I5 and I7 lines (blue, orange, and grey, respectively). (c) and (d) Volcano plots generated from 2 groups sample comparison tests for I3 vs. I5 (c) and I3 vs. I7 (d). Most statistically significant aldehydes as measured by p-value are shown labelled. See Supplemental Table S5 for detailed statistical analysis.

### 3.2. Polymorphisms in TAG lipases identified in the low rancid line are non-functional

The lipid and oxidative analyses suggest that these lines may have differences in TAG lipases. Our analysis identified a total of 44 lipase genes in pearl millet accessions in the International Pearl Millet Genome Sequence Consortium (IPMGSC) data. These lipases were divided into three major subfamilies based on sequence alignment and evolutionary relationships. Subfamily I and III categorised under PLD/PLC/esterase comprised of 12 and 18 genes, respectively (Figure 4a). Subfamily II comprised 14 genes, with a conserved LID domain and a catalytic triad, which are the characteristic features of TAG lipases. Subfamily II lipase gene and protein characteristics are summarized in Table S6.

**Figure 4.**
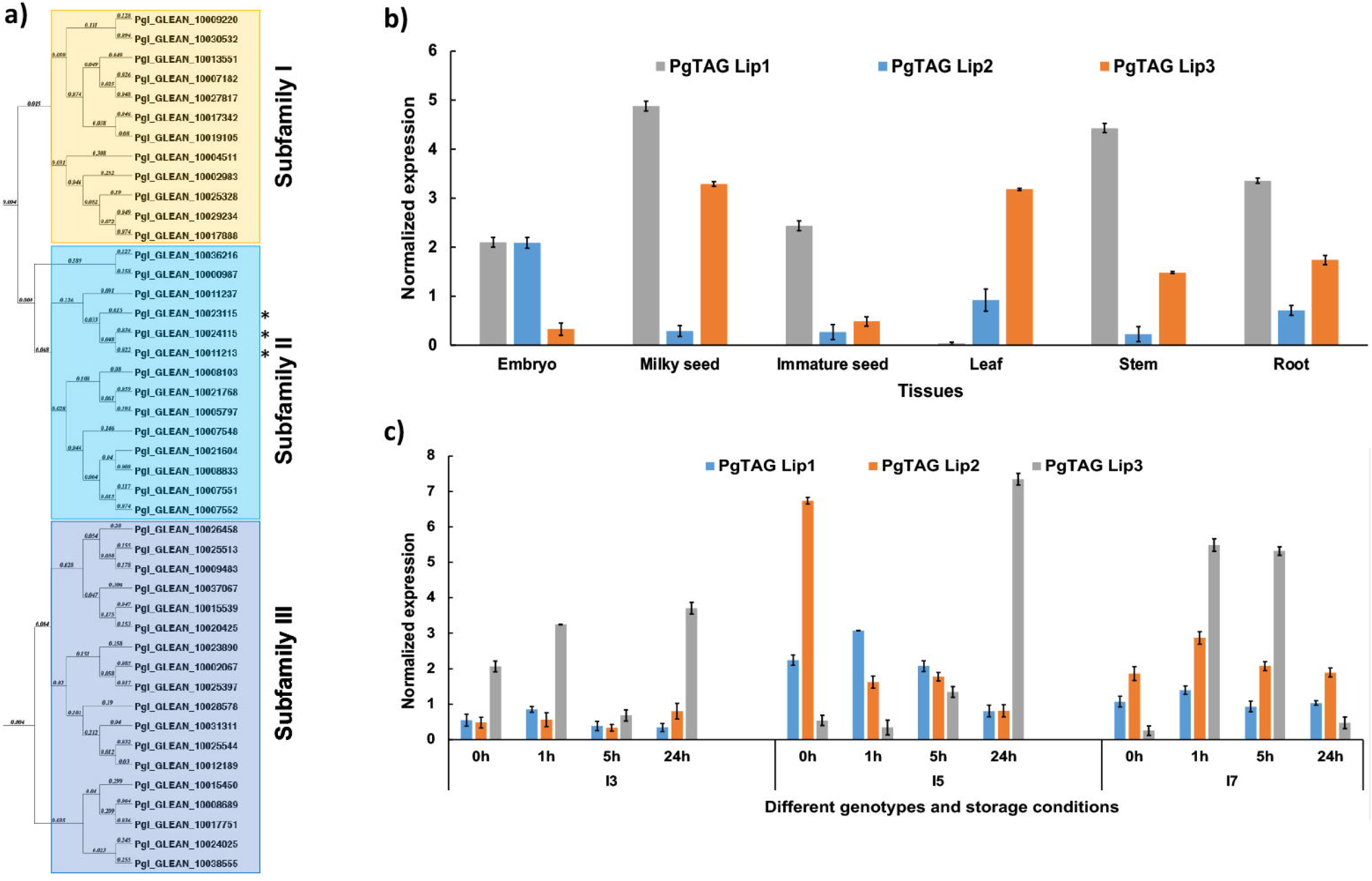
Phylogenetic relationships and expression analysis of the pearl millet lipases (a) The phylogenetic tree constructed using MacVector software by the NJ method for PgTAGLips. Asterisks (*) denotes the genes used for further analysis: Pgl_GLEAN_10023115 was renamed *PgTAGLip1*, Pgl_GLEAN_10024115 renamed *PgTAGLip2*, and Pgl_GLEAN_10011213 renamed *PgTAGLip3*. (b) Expression analysis of three selected PgTAGLips in the various developmental stages of pearl millet variety ICMB 9333. Data points represent the expression values, obtained after normalization against the reference genes *EUKARYOTIC INITIATION FACTOR4A* (*PgEIF4cα* and *MALATE DEHYDROGENASE (PgMDH*). Each data point represents mean of three biological replications with standard error (±SE) (representing mean of three technical replicates). (c) Expression of the selected PgTAG Lips in contrasting genotypes (I3, I5, and I7) under accelerated storage of milled flour. The X-axis represents the identities of flour samples stored at 0 h, 1 h, 5 h, 24 h. Y-axis indicates the normalized gene expression values against the reference genes *PgEIF4α* and *PgMDH*. Each data point represents mean of three biological replications with standard error (±SE) (representing mean of three technical replicates).

Based on the expression analyses of various plant tissues of the popular variety ICMB 9333 and 24 h accelerated storage of milled flour of the contrasting I3, I5, and I7 inbreds, three lipases were selected for further analyses. These lipases had higher similarity to the previously reported rice and Arabidopsis lipases (LOC_Os11g43510 and AT5G18640, respectively (Tiwari et al., 2016). The lipases Pgl_GLEAN_10023115, Pgl_GLEAN_10024115 and Pgl_GLEAN_10011213 were renamed PgTAGLip1, PgTAGLip2 and PgTAGLip3, respectively. *PgTAGLip2* had high expression in immature embryos of ICMB 9333 compared to other tissues such as milky seed, immature seed, leaf, stem and roots (Figure 4b). While *PgTAGLip1* expressed in most of the tissues, except leaf, with the highest expression at the milky seed stage, *PgTAGLip3* showed higher expression in milky seeds and leaves (Figure 4b). During 24 h of accelerated storage of flours from the contrasting inbred lines, there was a significant increase in the expression of all three tested genes in the flours of lines I5 and I7 compared to the flour from the line I3 (Figure 4c). Interestingly, the expression levels of *PgTAGLip1* and *PgTAGLip2* were considerably lower in I3 flour compared to the flours of I5 and I7 (>1.5 fold) during the first 24 h (Figure 4c), suggesting that these lipase genes may be responsible for the differences in TAG degradation between these lines (Figure 2).

Due to the differences in quantitative expression, the genetic sequences were analysed for structural polymorphisms across genotypes. Sanger sequencing confirmed that the full length coding regions of all three lipases carried a LID domain, an active site containing a serine GXSXG motif, and a catalytic triad (Figure S2). However, the *PgTAGLip1* gene sequences revealed polymorphisms among the lines I3, I5 and I7 (Figure 5a). To further confirm the correlation of these allelic variations with the observed and reported flour shelf life, additional genotypes (hybrids and inbred lines) based on a subset of the rancidity matrix were profiled for structural variations in *PgTAGLip1*. The *PgTAGLip1* gene in the I7 and high rancidity line 86M88 formed a functional protein encoded by a 1110 bp coding sequence, while the line I5 had a 807 bp coding region. Interestingly, the line I3 showed two transcript variants of Pg*TAGLip1* having fragment sizes of 528 bp and 276 bp (Table S6). These were named as LR1 and LR2, respectively. The LR1 transcript contained a stop codon within the LID domain, expected to result in a loss-of-function of this gene, while the LR2 variant formed a much smaller peptide lacking the LID domain/active site and is also presumed to be non-functional (Figure 5a). In addition to the line I3, other low rancidity lines Super Boss and RHB 177 also revealed mutations resulting in stop codons upstream of the LID domain at 420 bp and 285 bp, respectively (Figure 5a; Table S6). It was intriguing to note that in contrast to the *PgTAGLip2* transcript (1068 bp) in line I7, single base variations and deletions up to 6 bp were observed in both lines I3 and I5 causing premature stop codons (Figure 5b; Table S6). No sequence variations among the three tested genotypes were observed in *PgTAGLip3*. These results provide further evidence that the maintenance of TAG, lower accumulation of FFA, and lower amounts of aldehyde volatiles in the line I3 are associated with allelic variations within the *PgTAGLip1* and *PgTAGLip2* genes.

**Figure 5.**
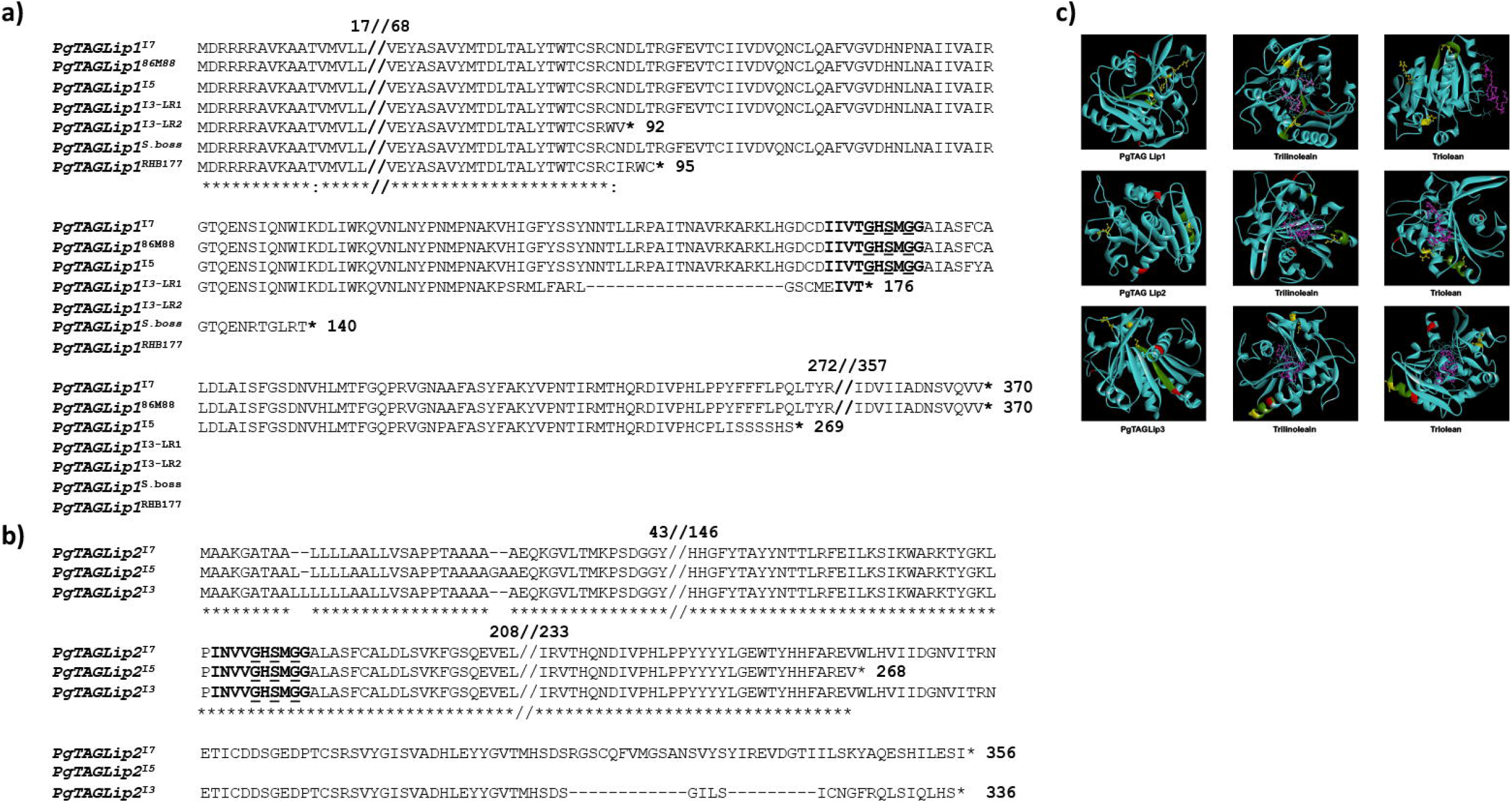
Structural variations of selected TAG lipases. (a) Comparative amino acid sequence alignment of Pgl_GLEAN_10023115 (PgTAGLip1) in high (I7, 86M88, I5) and low rancidity genotypes (I3, Super Boss, RHB177). The LID domain is in bold and the Lip serine active motif (GxSxG) is in bold and underlined. (b) Comparative amino acid sequence alignment of PgTAGLip2 in high rancid (I7, I5)and low rancid (I3) lines. The LIP domain is bold and Lip serine active motif (GxSxG) is bold and underlined. (c) Predicted protein models and docking affinities of three PgTAGLips (PgTAGLip1, PglTAGLip2, and PgTAGLip3) with Triolein and Trilinolein. The predicted substrate binding is shown in a ball and stick model, with dashed pink lines denoting the hydrogen bonds. The putative LID domain, GxSxG motif is in green, catalytic triad is in yellow with ball and stick, putative glycosylation sites in are in red, ligand ball and stick is in pink.

To further support our annotations, molecular docking was used to predict the affinity/specificity of these pearl millet TAG lipases to several triacylglyceride ligands (Table S7). *Rhizomucor meihei* lipase (RML) (PDB id: 3TGL) was used as a template to generate the three-dimensional models and shared a 32-55% peptide identity with the PgTAGLips. The modelled structure when superimposed with the template (PDB id: 3TGL) showed an overall root mean square deviation (RMSD) of 1.66 suggesting close structural similarities among the modelled and template structures. Geometrical aspects of the modelled structures revealed that >70% of the modelled secondary structure was favourable and based on the predicted Ramachandran plots, only 0.9% of residues fell in the disallowed region, thereby confirming the high quality of the modelled structures. Protein-ligand docking revealed that, PgTAGLip1, PgTAGLip2, and PgTAGLip3 had very high affinities/specificities for triglycerides containing unsaturated fatty acid residues, suggesting that these lipases have a predisposition for degrading TAG containing linoleic and oleic fatty acids in the pearl millet (Table S7; Figure 5c).

To determine if the PgTAGLip allelic variants were functional, they were assessed for complementation in a lipase-deficient yeast mutant. The yeast triple lipase mutant *Δtgl3Δtgl4Δtgl5* (ΔTGL) lacks all three endogenous lipases. *PgTAGLip1, PgTAGLip2*, and *PgTAGLip3* genes were cloned from the lines I3, I5 and I7 and transformed into ΔTGL. To indicate the germplasm source and genotypic identity of allelic variants (mutants), superscripts were added to the gene names, e.g. PgTAGLip1^I7^, PgTAGLip3^I3^, etc. The total amount of TAG from the transformed yeast cells was analysed by GC-FID. Yeast cells expressing full-length (non-truncated) PgTAGLip1, PgTAGLip2 and PgTAGLip3 accumulated less TAG (16.51, 18.72 and 14.65 μmol/g, respectively), when compared to the ΔTGL control (26.93 μmol/g) (Figure 6a). The truncated allelic variants PgTAGLip1^I3-LR1^, PgTAGLip1^I3-LR2^, PgTAGLip2^I3^ and PgTAGLip2^I5^ accumulated comparable TAG content to ΔTGL (Figure 6a) confirming that these truncations are non-functional.

**Figure 6.**
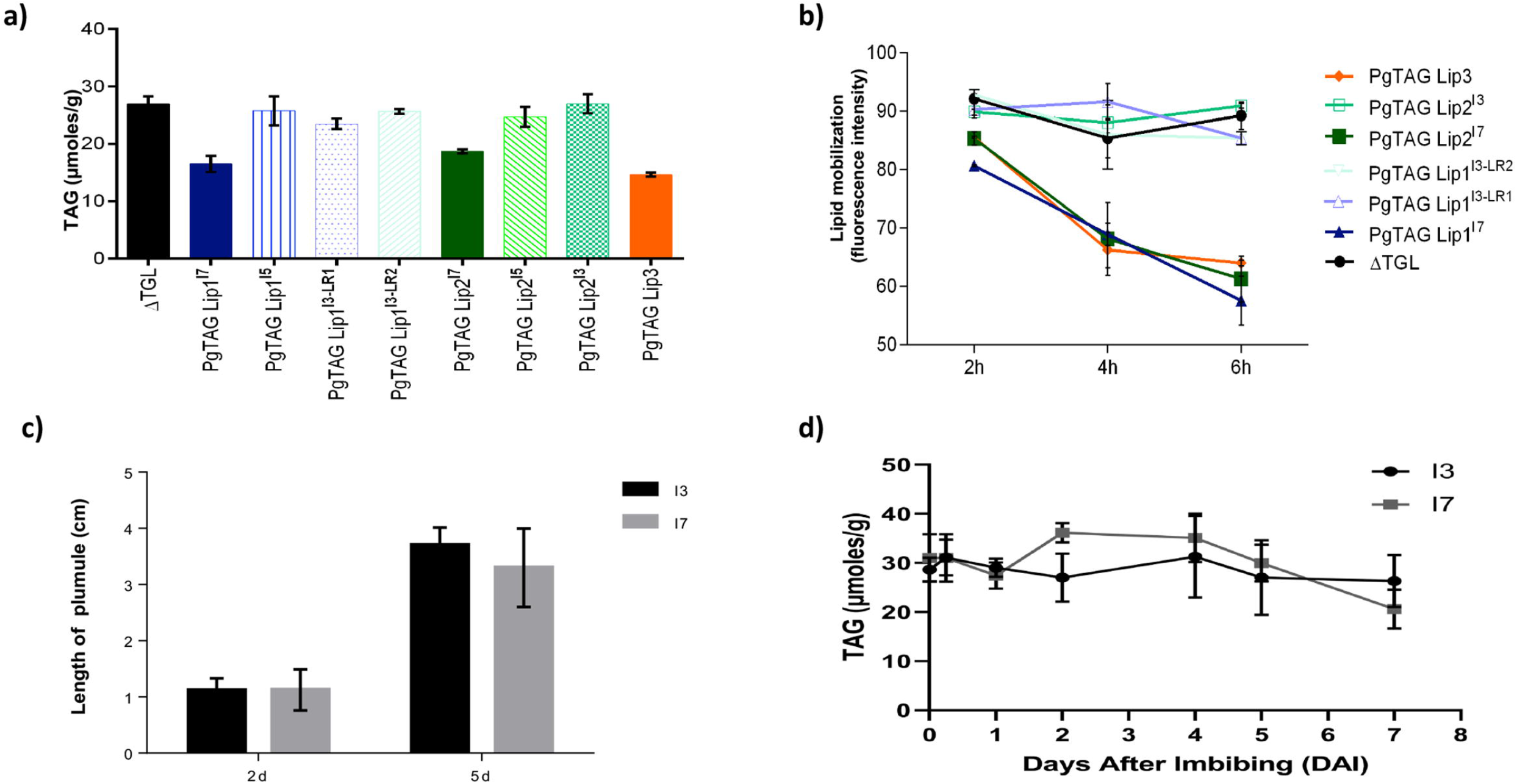
Functional validation in yeast expression system and seedling growth of the contrasting pearl millet seeds lines during germination (a) GC-MS TAG analysis of yeast cells expressing PM TAG lipase variants. Each data point represents mean of three biological replications with standard error (±SE) (b) *In vivo* mobilization of TAG reserve in yeast triple lipase mutant by TAG lipase variants from 2h-6h. Each data point represents mean of two biological replications with standard error (±SE). (c) No significant differences observed in germinated seedlings (d) TAG mobilization during germination and post-germinative growth. Each data point represents mean of three biological replications with standard error (±SE).

To corroborate the lower amounts of TAG from the truncated PgTAGLip variants, mean fluorescence intensity of yeast TAGs stained with BODIPY^493/503^ were analysed by flow cytometry. There was a significant decrease in fluorescence intensity in the populations of yeast cells (P3) expressing full-length lipase genes compared to the control (ΔTGL expressing empty vector). The yeast cells carrying non-truncated versions of PgTAGLip1, PgTAGLip3, and PgTAGLip2 cloned from the line I7 had lower intensities (64.25%, 58.10%, 54.15% respectively), reflecting TAG breakdown. In contrast, the cells expressing truncated variants PgTAGLip1 ^I3-LR1^, PgTAGLip1 ^I3-LR2^, PgTAGLip2 ^I3^, and PgTAGLip2 ^I5^ showed higher fluorescence intensities, viz. 82.65%, 84.75%, 82.55%, and 89.8%, respectively (Figure S3a). Hence, these results demonstrate that all the truncated allelic variants (PgTAG Lip1^I3-LR1^, PgTAG Lip1^I3-LR2^, PgTAG Lip2 ^I3^ and PgTAG Lip2 ^I5^) had reduced functionality.

### 3.3. TAG lipases with loss-of-function mutations do not mobilize TAG, which does not impact seedling establishment

The reduction in TAG content in yeast strongly suggested that all three non-truncated PgTAGLip exhibit lipolytic function. To demonstrate that deleterious mutations in the variants affect their function, time-dependent turnover of TAG was estimated during the yeast growth (up to 6 h) and measured using flow cytometry. Results indicated that the ΔTGL mutant and truncated variants PgTAGLip1^I3-LR1^, PgTAGLip1^I3-LR2^, PgTAGLip2^I3^ and PgTAGLip2^I5^ did not mobilize TAG products. In contrast, the non-truncated versions (cloned from the line I7) were significantly active in mobilizing TAG products during 2 h to 6 h time course (Figure 6b).

To determine if the observed differences in lipid degradation between the lines I3 and I7 had any effect on seedling establishment, plumule elongation and lipid mobilization were measured during germination. No differences were observed in post-germination seedling establishment or growth (Figure 6c), indicating that while the mutations in both candidate lipases may be useful for minimizing TAG hydrolysis, these enzymes are not essential for germination. Additionally, there were minimal differences in TAG mobilization between the lines I3 and I7 during germination (Figure 6d); most likely other TAG lipases are involved in mobilizing TAG for germination. Taken together, these results suggest that mutations in PgTAGLip1 and PgTAGLip2 contribute to higher amounts of FFA accumulating in mature seeds. Therefore, allelic variations in these genes may provide an opportunity to decrease the amount of FFA (and thus their oxidation), thereby alleviating the propensity of rancidity development in pearl millet flour.

## 4. Discussion

Considered as the “mother of all problems” in pearl millet, tackling the issue of flour rancidity is the most important intervention required for ensuring a sustainable demand for this nutri-cereal. While, stabilization techniques have been effective in minimizing rancidity to some extent, only large-scale processing would justify their economic feasibility which may not be practical in rural agricultural communities. The improved stability of shelf life following stringent treatments suggests that biological enzymatic-driven processes drive the generation of off-flavours volatiles in pearl millet flours. Thus, genetic improvements are a promising alternative to physical stabilization techniques for extending the storage life of flour.

### 4.1. Biochemical analyses suggest rancidity of pearl millet flour is due to lipid catabolism

A set of 12 pearl millet genotypes based on an earlier study (Majumdar et al., 2016) were investigated for identifying genotypes with contrasting susceptibility to rancidity. The change in AV, an indicator of esterified lipid hydrolysis and the presence of FFA over 10 days from seed grinding was used to categorise pearl millet lines. Contrasting inbred lines were categorised as low AV (I3), high AV (I5), and intermediate AV (I7) that were further confirmed by sensory panel analysis. Clear differences between the selected lines were observed in the amounts of TAG and FFA present in the seeds at the time of grinding through the accelerated-aging treatment, that were in broad agreement with the initial AV studies. Further analysis after 21 days of accelerated storage showed that aldehydes and other markers of fatty acids, primary and secondary oxidation (1-octen-3-ol and 2-pentyl furan) were the dominant discriminators in comparisons between the headspace volatiles of the flours from lines I3 vs I5 and I3 vs I7. Not only are these volatile chemicals key markers of oil oxidation (Ranklin and Mitchell, 2019), they are also associated with the development of rancidity in foods rich in polyunsaturated fats, including seeds and flours (Gorji et. al., 2019). Other, non-lipid related off-flavour metabolites such as 2-acetyl-1-pyrroline and apigenin have been identified in high moisture (30%) pearl millet flours within 15 h of grinding and wetting (Seitz el al., 1993) where both have been associated with a mousy acidic odour detected in millet flours. Mass spectral searches of the volatile chemistry profiles of the 21-day aged flours analysed in this study using the NIST-AMDIS application failed to detect these two compounds, thereby indicating that they were not principal factors in this material and that products of lipid oxidation were closely associated with flour rancidity.

These results indicate that differential hydrolysis of esterified lipids is linked to the release of FFA and is closely associated to the development of rancidity under accelerated-aging conditions. We presume that the amphipathic poly-unsaturated FFAs are more prone to oxidative degradation once released from lipid bodies and membranes resulting in the generation of short chain aldehydes and other secondary oxidation products characteristic of oil rancidity. Considering that the fatty acid profiles for the three lines were similar, these results indicate that the lines I5 and I7 had higher propensity for rancidity than the line I3 due to higher amounts of FFAs and the subsequent oxidation products. Furthermore, this direct association between TAG degradation and off-flavour volatiles suggest that there may be transcript or genomic differences in the TAG lipases between these pearl millet lines.

### 4.2 Non-functional mutations in PgTAGLip1 and PgTAGLip2 are associated with low rancidity

Three PgTAG lipases with close homology to those reported from *O. sativa* and *A. thaliana* lipases (Tiwari et al., 2016) were found to have differences in transcript levels and sequences between the three tested pearl millet lines. Expression analyses provided evidence that *PgTAGLip1, PgTAGLip2* and *PgTAGLip3* genes were expressed in stored flour, with the line I3 having the lowest level of expression of *PgTAGLip1* and *PgTAGLip2*, thereby suggesting a catalytic role in the degradation of triacylglycerols (TAGs) and FFA generation even after milling. Further insights into connecting these TAG lipase to the differential susceptibility to rancidity in the lines I3, I5 and I7 were revealed through sequence analysis where mutations were observed in the coding sequences of *PgTAGLip1* and *PgTAGLip2* of lines I3 and I5. Structural polymorphisms (SNPs and deletions) in one or both lipase genes *PgTAGLip1* and *PgTAGLip2* led to premature stop codons that terminated the *PgTAGLip* gene products before or close to the catalytic centre.

TAG lipases contain a catalytic serine-aspartate-histidine triad that is required for activity (Wada et al., 2009; Tiwari et al., 2016). A LID domain covering this active site is present in lipases and is critical for the interaction of lipase with the TAG substrate (El-Kouhen et al., 2005). Our study revealed, that in high rancid lines, *PgTAGLip1* in line I7 had an intact LID domain and catalytic triad, while *PgTAGLip1* had only the absence of histidine residue in the line I5 (Table S8). In contrast, the low rancid line I3 altogether lacked these active site motifs in this gene. Interestingly, while pre-mature truncations were apparent in *PgTAGLip2* gene in lines I3 and I5, the catalytic centre was intact in all three lines. While our study did not confirm the effect of truncations on the enzyme activity, the protein docking of various ligands (triglycerides) with both PgTAGLip1 and PgTAGLip2 predicted very high affinity/specificity for triolein and trilinolein substrates. These substrates are most likely some of the endogenous substrates, since oleic and linoleic fatty acids are major components of pearl millet flour (Zaplin *et al*., 2013). In addition to the *PgTAGLip1* polymorphisms uncovered in the line I3, detection of similar mutations in other low rancidity lines (Table S8) like Super Boss and RHB177, further imply that this TAG lipase is the root cause of rancidity in pearl millet. These findings are in line with previous reports where mutations in the serine residue in the GXSXG motif of *Pseudomonas aeruginosapatatin-like* phospholipase and yeast TAG Lipase 4 with an alanine residue almost demolished lipase activity (Kurat et al., 2006). Similarly, complete deletion of the GXSXG motif in mouse phospholipaseA2 (PLA2) almost eliminated its enzymatic activity (Wada et al., 2009). A yeast lipase-deficient system can be used to test the functionality of heterologous lipase genes (Bansal et al., 2021). Since the yeast storage lipids are structurally and functionally similar to seed storage lipids, they serve as ideal *in vivo* system to ascertain the role of candidate lipases in TAG hydrolysis (Jacquier et al., 2013). A comparison of the truncated lipases from the line I3 (*PgTAGLip1*^I3^ and *PgTAGLip2*^I3^) with the non-truncated lipases from the line I7 (*PgTAGLip1*^I7^ and *PgTAGLip2*^I7^) revealed that the lipase variants in line I3 were less functional in mobilizing yeast storage lipids. These results demonstrate that rendering PgTAGLip1 and/or PgTAGLip2 non-functional through mutations in the active site motifs can provide a mechanism for maintaining TAG levels and reduce off-flavour volatiles in the flour.

### 4.3. TAG Lipase polymorphisms can be used to develop pearl millet varieties with low susceptibility to rancidity

For the application of mutating TAG lipases to alleviate the rancidity of pearl millet flour and increase shelf life, physiological and agronomic effects must be minimal. Lipases are important in the catabolism of lipid reserves, especially during germination and post-germination seedling establishment (Eastmond, 2006). Our results show minimal changes to seedling establishment and TAG mobilization in the line I3 containing polymorphisms in *PgTAGLip1* and *PgTAGLip2*, thereby suggesting that these are not essential for germination or post-germination growth. While both these lipases may be useful for TAG hydrolysis, since the variants do not mobilize TAG, there are most likely functional redundancies among PgTAGLip enzymes.

## 5. Conclusion

We have identified specific polymorphisms in pearl millet lipases that can be used for breeding high yielding hybrid pearl millet varieties with prolonged flour shelf life. Our study also demonstrates strategies for future chemical mutagenesis or CRISPR-based approaches for creating loss-of-function mutations in *PgTAGLip1* or *PgTAGLip2* to mitigate rancidity in elite milled pearl millet germplasm. Increasing the storage life of flour from this nutritious grain offers tremendous opportunities for primary and secondary processing, creating markets and enhanced profits for smallholder farmers in South Asia and Sub-Saharan Africa.

## CRediT authorship contribution statement

Rasika Rajendra Aher: *Investigation, Validation;* Palakolanu Sudhakar Reddy: *Methodology, Original draft, Formal analysis;* Rupam Kumar Bhunia: *Methodology, Investigation, review & editing;* Kayla S. Flyckt: *Investigation, review & editing;* Aishwarya R Shankhapal: *Investigation, Validation;* Rabishankar Ojha: *Investigation;* John D. Everard: *Conceptualization, Methodology, Investigation, review & editing;* Laura L. Wayne: *Investigation, Formal analysis, review & editing;* Brian M. Ruddy: *Data curation, Visualization;* Benjamin Deonovic: *Data curation;* Shashi K. Gupta: *Methodology, review & editing;* Kiran K. Sharma: *Formal analysis, original draft, review & editing, Project administration;* Pooja Bhatnagar-Mathur: *Conceptualization, Funding acquisition, Methodology, Formal analysis, original draft, Supervision, Project administration*.

## Acknowledgements

RRA and RKB acknowledge fellowship support from the Department of Science and Technology in the form of INSPIRE Faculty award (DST/INSPIRE/03/2018/000417 & DST/INSPIRE/04/2017/000484 respectively). Authors also thank Central Instrumentation Facility (CIF)-NABI for providing key instruments used during the study. Thanks to Joseph D. Shambaugh for assistance with workflow development on the Genedata Expressionist software platform. Authors thank Professor Günther Daum, Graz University of Technology for kindly providing the yeast lipase-mutant strain. PBM acknowledges the financial support from the CGIAR Research Program on Grain Legumes and Dryland Cereals (CRP-GLDC) supported by CGIAR Fund Donors

**<u>Appendix A,B & C. Supplementary data</u>**

