## Supplementary data for "Loss-of-function mutations in novel triacylglycerol lipase genes are associated with low rancidity in pearl millet flour"

Supplementary Materials

### Appendix A. Supplementary methods

#### 1. Sensory evaluation of flours under accelerated storage

A preliminary sensory panel of basic attributes to profile the aroma, flavour, and taste of flours from the selected inbred lines was carried out at ICRISAT (Table S3). The samples were presented to a panel of six assessors randomly in replicates of 5 at regular intervals during accelerated storage (up to 21 days), to characterize the differences in aroma and taste. The assessors were advised to not differentiate by colour. The responses were summarized based on the frequency of descriptors. Ground coffee powder was used to clear the olfactory sensors between sample evaluations.

#### 2. Lipid quantitation by TLC/GC-MS

Total lipids from pearl millet flour and yeast cells were extracted according to Sinha et al. (2020) and were separated through thin layer chromatography (TLC). For GC-MS analysis, FAMEs of TAG and FFAs were prepared from the individual lipids using 4% (v/v) sulfuric acid in methanol along with 0.2 mL toluene and 10 μg of heptadecanoic acid (17:0, Sigma, USA) as an internal standard. FAMEs were quantified using Agilent Technologies Model 7890B gas chromatograph equipped with 7000 GC/MS triple quad detector and a non-polar DB-5 capillary column (J&W 123–5533, 30 m x 320 μm × 1 μm) according to Sinha et al. (2020). The FA concentrations were determined using the known concentration of 17:0, represented as a percent of dry weight (%DW) and μmoles/g dry weight (μmol/g DW).

#### 3. Statistical Modeling

Separate models were fit for TAG and FFA. Several models were fit all of which included line as a fixed effect to allow for inference. Other factors considered for inclusion in the model are time (measured in days since start of experiment; 0, 7, 14, and 21 days) and an indicator from which experiment the data are from (i.e. collection: Corteva-2019, Corteva-2021, and NABI-2021). A

random effect was included to account for the correlation among technical replicates. Models including all combinations of the fixed effects and interactions of the fixed effects were fit. The FFA values in two of the collections (Corteva-2019 and Corteva-2021) exhibited left censoring of several values due to lower levels of detection. These values were included in the analysis as censored values with the censoring threshold set to the minimum of the non-censored observations from these two collections.

The models were fit using the brms software package fit using the brms software package (Bürkner, 2017) in the R programming language (R Core Team, 2021). Models were compared to select a model which balances parsimony and fit to the data by comparing and selecting the model with the lowest leave-one-out information criteria (Vehtari et al., 2017). The models selected for both TAG and FFA analysis included the time component (allowing for changing TAG and FFA levels over time) as well as an interaction between time and line type (allowing varying trends between each line). The model selected for TAG additionally included collection as a covariate (allowing for varying initial TAG levels), while the FFA model included collection as a covariate as well as the interactions between collection with time and line (allowing for varying trends between the lines across the three collections).

#### 4. Solid Phase Micro-Extraction-GC-MS parameters & data analysis

For all samples and controls, the headspace vials were agitated (250 rpm; 70˚C) for 5 min prior to SPME (Supelco 57298-U; Gray) headspace sampling for 20 min. With the inlet (0.75 mm ID liner; Restek) set at 250 °C in a splitless mode, the fibre was exposed to start the cycle. After 5 min in the inlet the split vent was opened (30mL/min) and was retained until the next cycle began. Chromatographic separation was achieved on an Agilent Ultra-1 column (50m x 0.32, 0.52u film thickness; helium carrier gas; 1.1 mL / min constant). The column oven was maintained at 40 °C for the first 5 min after sample introduction and was then ramped (12 °C /min) to 240°C, with a 1 min hold, before ramping (30 °C/min) to 325 °C, which was held for 4 min prior to returning to the starting conditions.

The MS data files were pre-processed and analyzed with Genedata Expressionist software tools. Workflow included: nominal mass integration (centroids integrated to nominal mass value

peaks), chromatogram blank subtraction, chemical noise subtraction, extracted ion chromatogram alignment, retention indexing (ethyl ester; C2-C14), chromatogram nominal mass peak detection and component detection (library matching; NIST v 2.2). Statistical workup was performed in Genedata Analyst on the weight normalized intensity of the most abundant ion representing the deconvoluted EI fragment groups. K groups and analysis of variance (ANOVA) were performed between all three lines and 2 groups (group medians) were performed for comparison between the I3 line and the I5 or I7 lines portrayed as volcano plots. Principal component analysis (PCA) modeling was performed by PLS_Toolbox (Eigenvector Research, Inc.) within MATLAB and featured autoscaling to accommodate the magnitude in the MS signal dynamic range.

### Appendix B. Supplementary Figures

**Figure S1.** TAG and FFA levels in milled flour under accelerated-ageing conditions compared across three experiments. TAG and FFA levels measured by HPLC-ELSD in two datasets (2019 and 2021; same data as Figure 2c) compared to a third dataset (NABI-2021) measured by TLC/GC-MS. Log-linear hierarchical mixed models were utilized to assess the differences in TAG and FFA levels of these three inbred lines. Points represent observed data, lines represent posterior mean of the predictive distribution, and the ribbon corresponds to the 95% credible interval of the posterior mean predictive distribution.

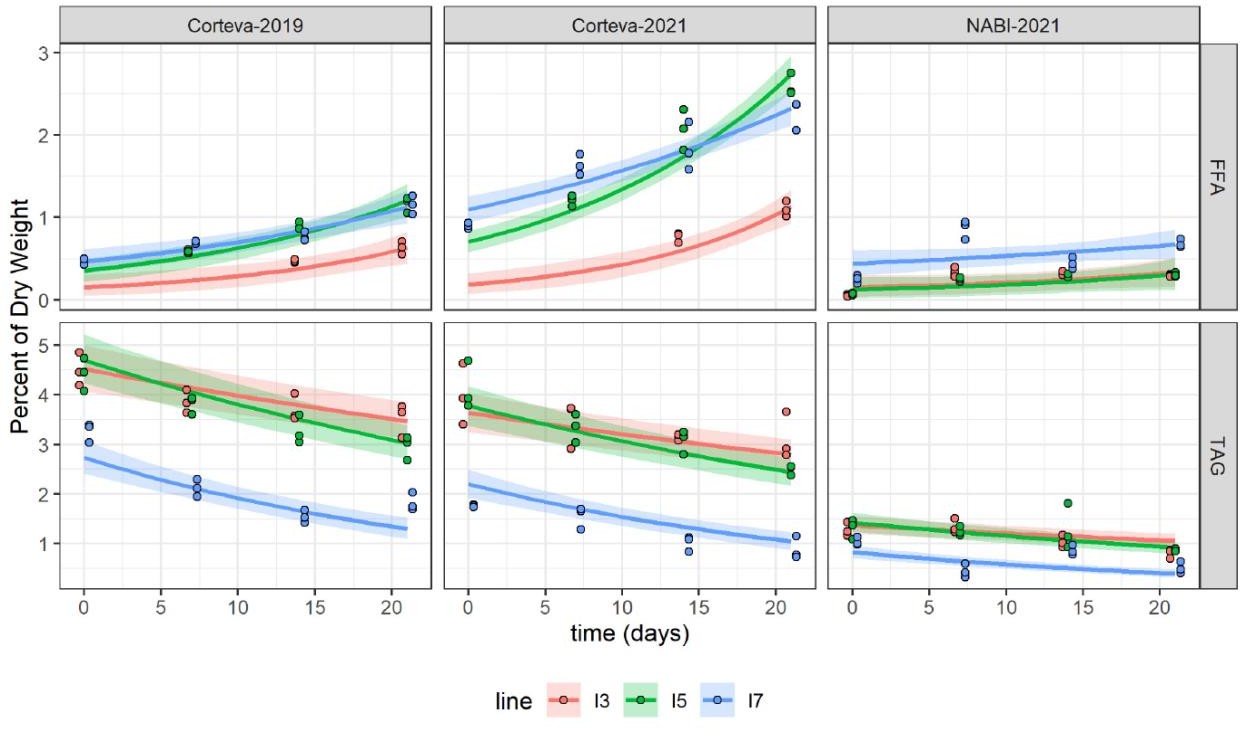

**Figure S2.** Sequence variations of selected TAG lipases. Comparative amino acid sequence alignment of three PgTAGLips, PgTAGLip1, PgTAGLip2 and PgTAGLip3 using the Clustal W program. The amino acids Ser (S), Asp (D) and His (H) residues forming the catalytic triad of TAG Lips are indicated with # symbol. The LID domain in bold letters carries the underlined lipase serine active motif (GxSxG). The putative N-glycosylation sites are indicated in red.

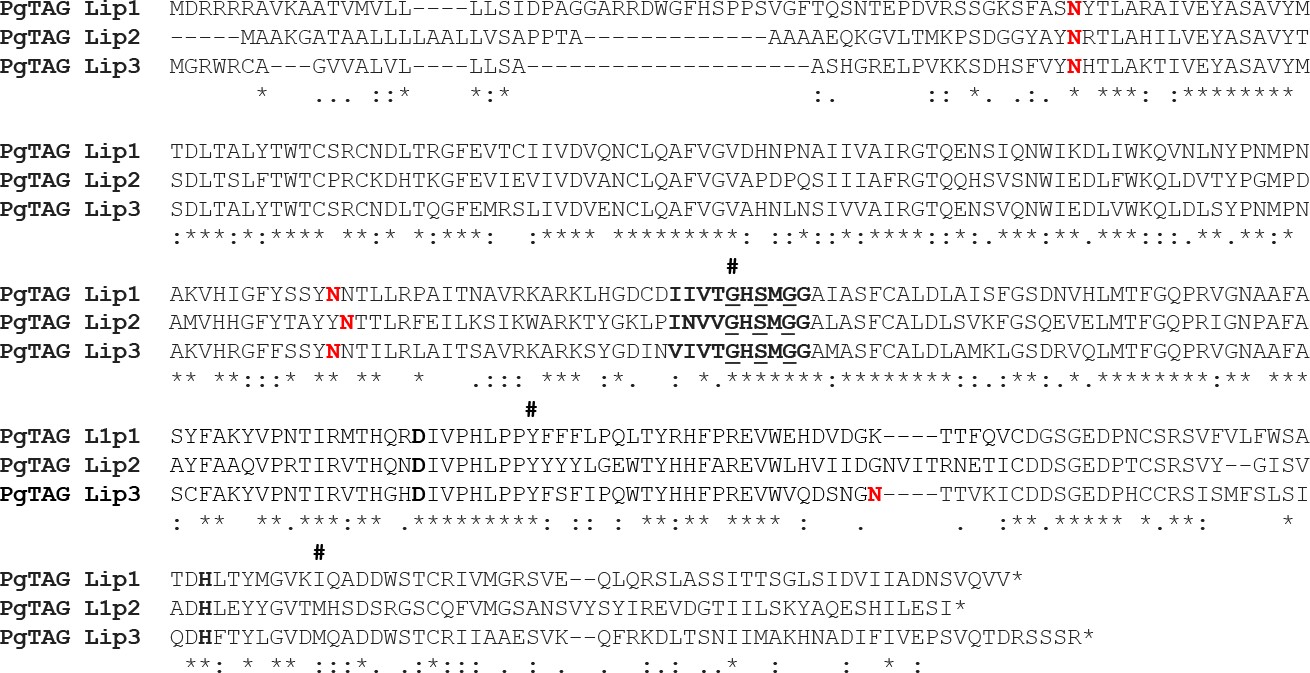

**Figure S3.** (a) Quantification of intracellular yeast neutral lipid bodies using flow cytometry. The X-axis represent fluorescein isothiocyanate-A (FITC-A) mean fluorescence intensity, where Y-axis represents cell count showing fluorescence. Increased fluorescence indicates greater numbers of lipid bodies. (b) Expression of indicated TAG lipase variants in ΔTGL yeast strain alters TAG content accumulation identified by FACS. Each data point represents mean of two biological replications with standard error.

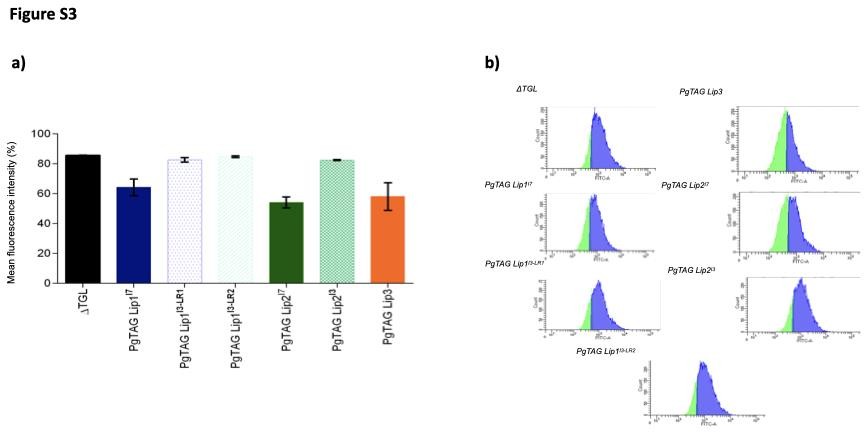

### Appendix C. Supplementary Tables

**Table S1:** Identities and codes of pearl millet genotypes used in this study.

| **Genotypes** | **Codes** |
| --- | --- |
| ICMB-843 | I1 |
| ICMB-88004 | I2 |
| ICMB-95222 | I3 |
| ICMB-81 | I4 |
| ICMB-89111 | I5 |
| ICMB-842 | I6 |
| ICMB- 863 | I7 |
| ICMB-98222 | I8 |
| IP5931 | P13 |
| IP 13840 | P14 |
| IP 22419 | P19 |
| IP 6099 | P20 |

#### Additional genotypes used in molecular analysis based on Goswami et al., 2020.

**Rancidity Potential**

**Genotypes**

86M88	High

Super Boss	Low

RHB 177	Low

**Table S2**. List of the primers used for full-length gene cloning, qRT-PCR and yeast expression analysis.

| **S.No** | **Gene** | **Primers** | **Remarks** |
| --- | --- | --- | --- |
| 1 | PgTAG Lip1 _F | ATGGATAGGCGGAGACGCGCGGT | Full-length gene cloning |
| 2 | PgTAG Lip1_R | TCAAACGACCTGGACGCTATTGTC |  |
| 3 | PgTAG Lip2_F | ATGGCTGCGAAGGGCGCGAC |  |
| 4 | PgTAG Lip2_R | TCATATGGATTCTAGGATATG |  |
| 5 | PgTAG Lip3_F | ATGGGGAGATGGAGGTGCGC |  |
| 6 | PgTAG Lip3_R | CTATCTAGAACTGCTCCGAT |  |
| 7 | PgTAG Lip2_qF | GGGCTTTGAGGTGATTGAGGT | Expression analysis |
| 8 | PgTAG Lip2_qR | ACACTGTGCTGTTGAGTCCC |  |
| 9 | PgTAG Lip3_qF | GCTGCAGGTCCATCTCCATG |  |
| 10 | PgTAG Lip3_qR | TGCTGGTGAGATCCTTTCGG |  |
| 11 | PgTAG Lip1 _qF | GCGTGTTTGTGCTGTTCTGG |  |
| 12 | PgTAG Lip1_qR | TTGCCCCATGACGATTCTGC |  |
| 13 | PgEIF4A_qF | ATCGTGAGCTTTACATCCATCG |  |
| 14 | PgEIF4A_qR | TATCCCTCAGGATACGGATGTC |  |
| 15 | PgMDH_qF | AGAAGGCGCTTGCTTACTCAT |  |
| 16 | PgMDH_qF | CAGTTCTGGGTGAGGGAATCT |  |
| 17 | PgTAGLip1_I7YF | ATTAGGTACCATGGATAGGCGGAGACGCGC | Yeast expression |
| 18 | PgTAGLip1_I7YR | CGCAGAGCTCTCAAACGACCTGGACGCTAT |  |
| 19 | PgTAGLip1_I5YF | ATTAGGTACCATGGATAGGCGGAGACGCGC |  |
| 20 | PgTAGLip1_I5YR | CGCAGAGCTCTCAGCTGTGGGAGGAAGAA |  |
| 21 | PgTAGLip1_I3LR1_YF | ATTAGGTACCATGGATAGGCGGAGACGCGC |  |
| 22 | PgTAGLip1_I3LR1_YR | CGCAGAGCTCTTATGTCACAATCTCCATG |  |
| 23 | PgTAGLip1_I3LR2_YF | ATTAGGTACCATGGATAGGCGGAGACGCGC |  |
| 24 | PgTAGLip1_I3LR2_YR | CGCAGAGCTCTCAAACCCATCTTGAGCAT |  |
| 25 | PgTAG Lip2_I7_YF | ATTAGGTACCATGGCTGCGAAGGGCGCG |  |
| 26 | PgTAG Lip2_I7_YR | CGCAGAGCTCTCATATGGATTCTAGGAT |  |
| 27 | PgTAG Lip2_I5_YF | ATATAAGCTTATGGATAGGCGGAGACGCGC |  |
| 28 | PgTAG Lip2_I5_YR | AGATGGATCCTCCTAAACCTCTCTAGCAAA |  |
| 29 | PgTAG Lip3_YF | ATTAGGTACCATGGGGAGATGGAGGTGC |  |
| 30 | PgTAG Lip3_YR | CGCAGAGCTCCTATCTAGAACTGCTCC |  |

**Table S3.** Sensory evaluations of the milled flour from lines i3, i5, and i7 during 21 days of storage under accelerated aging conditions.

| **Days of storage** | **Odour** | | |  | **Taste** | | |
| --- | --- | --- | --- | --- | --- | --- | --- |
|  | **I3** | **I5** | **I7** |  | **I3** | **I5** | **I7** |
| 0 | Dry flour | Cooked rice | Pungent |  | Spoiled rice | Bitter gourd | Bitter |
|  | Dry flour | Fresh dry flour | Light acid smell |  | Bitter | Light bitter | Slightly bitter |
|  | Wheat dough | Dry flour | Slightly flour like smell |  | Slightly sweet | Flour like | Bitter |
| 1 | Sweet | Dry flour | Light pungent |  | Bitter | Bitter | Bitter |
|  | Mild flour | Wet mud | Stale dough |  | Sweet wheat flour |  | Slightly bitter |
|  | Fermented dough | Sweet | Pungent |  | Tasteless | Slightly bitter | Bitter |
| 7 | Fermented batter | Light gym socks | Stored rice flour |  | Sour | Bitter | Camphor |
|  | Sweet | Fermented dough | Foul |  | Bitter | Bitter | Bitter |
|  | No odour | Pungent | Damp-dough |  | Slight bitter | Bitter corn | Bitter |
| 14 | Strong wheat flour | Stored rice | Gym socks |  | Bitter as bitter gourd | Slightly bitter | Chalk powder |
|  | No pungency | Slightly stale wheat | Strong bad smell |  | No taste | Bitter | Mud like |
|  | Sweet smell | Bad smell | Woody, wet soil, tree bark smell |  | Sour | Bitter | Bitter |
| 21 | General flour smell | Foul | Stored rice bags |  | Bitter and sour | Stored cornflakes | Stored wheat flour |
|  | Very light pungent | Very light pungent | Light pungent |  | No Taste | Very bitter | Spoiled flour taste |
|  | Strong flour | Pungent | Damp/woody |  | Light bitter | Bitter | Mud like |

-*#*	--Internal Use---

**Table S4.** Pairwise contrasts of line TAG (left) or FFA (right) values shown in Figure 2c. Pairwise median posterior estimates of differences between (log) TAG values for each line pair at each time point along with 95% highest probability density credible interval.

**TAG**	**FFA**

**Days contrast Estimate lower.HPD upper.HPD sig	Days	contrast Estimate lower.HPD upper.HPD sig**

|  | I3 - I5 | -0.026 | -0.118 | 0.069 | - |  |  | I3 - I5 | -1.302 | -2.05 | -0.685 | * |
| --- | --- | --- | --- | --- | --- | --- | --- | --- | --- | --- | --- | --- |
| **0** | I3 - I7 | 0.564 | 0.44 | 0.692 | * |  | **0** | I3 - I7 | -1.722 | -2.43 | -1.088 | * |
|  | I5 - I7 | 0.59 | 0.469 | 0.718 | * |  |  | I5 - I7 | -0.421 | -0.646 | -0.193 | * |
|  | I3 - I5 | 0.04 | -0.042 | 0.115 | - |  |  | I3 - I5 | -1.146 | -1.619 | -0.724 | * |
| **7** | I3 - I7 | 0.732 | 0.631 | 0.833 | * |  | **7** | I3 - I7 | -1.377 | -1.857 | -0.974 | * |
|  | I5 - I7 | 0.693 | 0.59 | 0.791 | * |  |  | I5 - I7 | -0.232 | -0.385 | -0.083 | * |
|  | I3 - I5 | 0.106 | 0.022 | 0.191 | * |  |  | I3 - I5 | -0.991 | -1.252 | -0.771 | * |
| **14** | I3 - I7 | 0.9 | 0.767 | 1.028 | * |  | **14** | I3 - I7 | -1.033 | -1.296 | -0.813 | * |
|  | I5 - I7 | 0.795 | 0.667 | 0.926 | * |  |  | I5 - I7 | -0.042 | -0.144 | 0.058 | - |
|  | I3 - I5 | 0.171 | 0.058 | 0.276 | * |  |  | I3 - I5 | -0.832 | -1.022 | -0.65 | * |
| **21** | I3 - I7 | 1.067 | 0.887 | 1.269 | * |  | **21** | I3 - I7 | -0.682 | -0.877 | -0.494 | * |
|  | I5 - I7 | 0.896 | 0.71 | 1.094 | * |  |  | I5 - I7 | 0.148 | 0.033 | 0.263 | * |

*#*---Internal Use---

**Table S5.** Statistical analysis for the SPME-EI-GC data shown in Figure 3. Analysis of variance (ANOVA) from K Groups (group medians) sample comparison tests for the four most abundant aldehydes in the mass spectral variable set. *p* -values and adjusted *p* -values (Benjamini Hochburg (BH Q- Value) are shown.

| **Analysis of Variance** | ***p* -Value** | **BH Q-Value** |
| --- | --- | --- |
| hexanal | 6.00E-07 | 2.00E-03 |
| benzeneacetaldehyde | 2.20E-06 | 3.60E-03 |
| octanal | 1.30E-05 | 1.30E-02 |
| nonanal | 9.00E-05 | 3.00E-02 |
| heptanal | 2.60E-04 | 4.90E-02 |
| pentanal | 4.80E-04 | 6.70E-02 |
| decanal | 7.50E-01 | 9.40E-01 |

-*#*	--Internal Use---

**Table S6.** List of the subfamily II lipase genes identified in this study and genomic and protein characteristics. Common properties such as open reading frame (ORF), amino acid translations, molecular weight (kDa) and isoelectric point (pI) of each pearl millet TAG Lipase (PgTAGLip) were calculated using the MacVector software (V17.1). Genes in bold were named and further characterised.

| **Gene Accession No.** | **Putative functional annotation** | **Lip domain/GXSXG motif** | **Major regions/domain** | **Nucleotide** | | | **Protein** | | | |
| --- | --- | --- | --- | --- | --- | --- | --- | --- | --- | --- |
|  |  |  |  | **ORF size** | **Chromosome** | **Chromosome location** | **Amino acids** | **MW** | **pI** | **Subcellular localization*** |
| **Pgl_GLEAN_10007548** | **phospholipase A1-II 3** | **ITITGHSLGG (106 - 115)/GXSXG** | **lipase class 3** | **792** | **chr6** | **104623040:104626472** | **263** | **28** | **5.04** | **chloroplast** |
| **Pgl_GLEAN_10007551** | **phospholipase A1-II 2-like** | **ITVVGHSLGA (160 - 169)/GXSXG** | **lipase class 3** | **1059** | **chr6** | **104679622:104682716** | **352** | **39.2** | **5.7** | **cytosol** |
| **Pgl_GLEAN_10007552** | **phospholipase A1-II 1** | **ITITGHSLGA (219 - 228)/GXSXG** | **PPR_2/lipase class 3** | **1185** | **chr6** | **104685154:104687393** | **394** | **44.07** | **6.33** | **mitochondria** |
| **Pgl_GLEAN_10021604** | **phospholipase A1-II 7-like** | **ITITGHSLGA (11 - 20)/GXSXG** | **lipase class 3** | **582** | **chr1** | **258491452:258492033** | **193** | **20.84** | **5.62** | **endoplasmic reticulum** |
| **Pgl_GLEAN_10008833** | **Phospholipase A1-IIgamma** | **ITVTGHSLGA (122 - 131)/GXSXG** | **lipase class 3** | **588** | **chr1** | **233144775:233146216** | **195** | **21.56** | **5** | **chloroplast** |
| **Pgl_GLEAN_10000987** | **uncharacterized protein** |  | **Lipase_3/catalytic triad** | **672** | **chr6** | **chr6:107492484:107494611** | **223** | **24.46** | **9.54** | **nucleus** |
| **Pgl_GLEAN_10008103** | **phospholipase A1-Ibeta2** | **ITVVGHSLGA (142 - 151)** | **triacylglycerol lipase/PLN02802/catalytic triad** | **981** | **chr3** | **34308672:34309652** | **326** | **35.33** | **8.68** | **chloroplast** |
| **Pgl_GLEAN_10021768** | **phospholipase A1-Ibeta2/** | **ITIVGHSLGA (153 - 162)** | **triacylglycerol lipase/PLN02802/catalytic triad** | **-** | **chr5** | **99694060:99695920** | **-** | **-** | **-** | **chloroplast** |
| **Pgl_GLEAN_10005797** | **phosphoinositide phospholipase C 2-like** |  | **PA2_HIS** | **354** | **chr2** | **chr2:18365615:18367043** | **117** | **12.72** | **5.48** | **extracellular** |
| **Pgl_GLEAN_10011237** | **lipase like** |  | **Lipase_3/catalytic triad** | **1029** | **chr1** | **163462080:163464746** | **342** | **38.24** | **5.84** | **extracellular** |
| **Pgl_GLEAN_10024115** | **lipase like** | **INVVGHSMGG(178 - 187)** | **Lipase_3/catalytic triad** | **1068** | **chr1** | **7199095:7202565** | **355** | **39.29** | **5.94** | **extracellular** |
| **Pgl_GLEAN_10023115** | **lipase-like** | **IIVTGHSMGG (192-201)** | **Lipase_3/catalytic triad** | **1110** | **-** | **-** | **370** | **34.88** | **8.77** | **vacuole** |
| **Pgl_GLEAN_10011213** | **lipase like** | **VIVTGHSMGG (169 - 178)** | **Lipase_3/catalytic triad** | **1056** | **chr7** | **1341282:1344315** | **351** | **39.29** | **7.63** | **chloroplast** |
| **Pgl_GLEAN_10036216** | **monoglyceride lipase** | **GXSXG (118-122)** | **hydrolase; alpha/beta fold family protein** | **909** | **chr6** | **234243736:234247360** | **302** | **33.33** | **6.9** | **cytosol** |

-*#*	--Internal Use---

**Table S7.** Proteins encoded by selected subfamily II lipases in pearl millet, their docking score, and observed hydrogen bonds between various docked substrates (triglycerides). The lower the score, the greater the preference for that substrate.

| **Gene Name** | **pNPA** | **Triacetin** | **Triarachidin** | **Tricaprin** | **Trilinolein** | **Triolein** | **Tripalmitin** | **Tristrearin** |
| --- | --- | --- | --- | --- | --- | --- | --- | --- |
| Pgl_GLEAN_10023115 (*PgTAGLip1* ) | -3.45 | -2.72 | -11 | 0 | -1.9 | -4.5 | -2.5 | -2 |
| Pgl_GLEAN_10024115(*PgTAGLip2* ) | -3.14 | -2.2 | -10.79 | -1.2 | -8.2 | -4.1 | -0.6 | -0.9 |
| Pgl_GLEAN_10011213(*PgTAGLip3* ) | -2.9 | -2.3 | -11.4 | -2.5 | -8.7 | -3.5 | -0.5 | -0.9 |
| Pgl_GLEAN_10008103 | -3.45 | -3.5 | -10.8 | -5 | -7.2 | -3.8 | -0.5 | -0.28 |
| Pgl_GLEAN_10007551 | -6.64 | -4.84 | -7.64 | -2.72 | -3.42 | -2.7 | -4.44 | -3.83 |
| Pgl_GLEAN_10021604 | -5.95 | -4.35 | -9.39 | -1.35 | -3.23 | -1.1 | -3.93 | -5.32 |
| Pgl_GLEAN_10007548 | -6.85 | -5.15 | -5.17 | -0.3 | -3.69 | -0.86 | -4.85 | -5.99 |
| Pgl_GLEAN_10007552 | -6.11 | -3.6 | -7.97 | -0.61 | -3.04 | -0.71 | -4.19 | -3.6 |
| Pgl_GLEAN_10008833 | -6.27 | -4.67 | -7.82 | -1.09 | -2.11 | -0.66 | -3.07 | -3.46 |
| Pgl_GLEAN_10011237 | -5.9 | -4.28 | 8.73 | -1.04 | 0.2 | -0.82 | 1.26 | 5.84 |
| Pgl_GLEAN_10036216 | -6.54 | -4.84 | 7.24 | -2.72 | 3.22 | -2.7 | 4.44 | 3.83 |

**Table S8.** Allelic variations found in of high (I7, 86M88, I5) and low rancid lines (I3 (LR1, LR2, Super Boss, RHB177) of Pg*TAGLip1* and Pg*TAGLip2.*

| **Gene** | **Genotype** | **ORF size (Nucleotides)** | **Protein size (Amino acids)** | **Lid domain/GXSSG motif/Catalytic triad** |
| --- | --- | --- | --- | --- |
| *PgTAG Lip1* | I7 | 1110 | 370 | Present |
|  | 86M88 | 1110 | 370 | Present |
|  | I5 | 807 | 269 | Histidine residue absent |
|  | I3 LR1 | 528 | 176 | Absent |
|  | I3 LR1 | 276 | 92 | Absent |
|  | Super boss | 420 | 140 | Absent |
|  | RHB177 | 285 | 95 | Absent |
|  | RHB223 | 276 | 92 | Absent |

| **Gene** | **Genotype** | **ORF** | **Protein** | **LID domain/GXSSG motif/Catalytic triad** |
| --- | --- | --- | --- | --- |
| *PgTAG Lip2* | I7 | 1068 | 356 | Present |
|  | I5 | 804 | 268 | Present |
|  | I3 | 1008 | 336 | Present |
